# Genomic tessellations

**DOI:** 10.64898/2026.11.13.766490

**Authors:** Barış Ekim, Haoyu Cheng

## Abstract

Periodicity is ubiquitous in genomic data. Assembled sequences are dense with homopolymers, microsatellites, and large satellite arrays, which contribute to phenotype, variation, and disease. Third-generation sequencing technologies, such as Oxford Nanopore (ONT) and PacBio HiFi, has enabled longer and increasingly accurate reads, but have brought with them systematic error biases in repetitive regions, where insertions and deletions can distort array lengths and mimic or obscure biological variation. This periodicity is not benign: local repetitiveness makes sequence alignment ambiguous and genome assembly error-prone. In sequence analysis, these two manifestations of periodicity are simultaneously at play, each compounding the challenges of the other; a unified representation of repetitive and non-repetitive sequence is necessary to account for their combined effects consistently.

We propose a paradigm under which (either repetitive and non-repetitive) DNA is split into its *identity* (its irreducible components) and its *geometry* (how its components are realized). A *genomic tessellation τ* is a canonical factorization of any sequence *S* of length *n* over the DNA alphabet into a sequence of *tiles* partitioning *S*, where each *t* in *τ*_*S*_ is described as a quasi-periodic run of a canonical unit *identity u*_*t*_ with *geometry* consisting of span *s*_*t*_, phase ϕ_*t*_, and a set of *exceptions X*_*t*_. We show that, for a prescribed error bound *k ≥* 1 and fixed maximum unit length *P, S* can be tessellated in *O*(*n*) time such that every tile has a unique identity whose periodic expansion differs from the tiled sequence by at most *k* edits, and that tiles sharing an identity are related by a bounded number of geometric adjustments. We introduce the *tessellated Burrows-Wheeler transform* (tBWT), which indexes *ω*_*S*_ by concatenating, over all tiles in order, the identity *u*_*t*_ of each *t*. For a tessellated query *q* of *m* tiles, the tBWT counts the repeat-agnostic occurrences of *q* in *O*(*m*) time, and for sampling interval *r* locates its *h* occurrences in *τ* (*S*) in *O*(*m* + *hr*) time.

We demonstrate that tessellations avoid the well-known speed-accuracy tradeoff and outperform state-of-the-art long-read mappers in repeat-dense regions in both speed and accuracy. On a 20× simulated set of both ONT and HiFi reads on the HG002 maternal X chromosome with synthetically-expanded repeat arrays, locating and chaining exact tile identity matches obtained from the tBWT is up to 57× and 83× faster, respectively, than the state-of-the-art mappers `minimap2` and `Winnowmap2`, with comparable accuracy. Furthermore, by abstracting away differences in geometry, tessellations of a set of input sequences onto itself (an *autotessellation*) reveals homology otherwise obscured by array-length variation and local sequence divergence. A geometry-only polishing of an an ONT-only assembly of the HG002 maternal X chromosome with ∼50× PacBio HiFi reads improves assembly QV from 45.4 to 60.6 in under a minute using less than 10GB of memory. Tessellated-read evidence over the autotessellation of this assembly organizes 167 mitochondrial contigs totaling ∼4Mbp into a single family with a ∼16.5kbp representative, which aligns to the HG002 Q100 mitochondrial genome with 100% identity. Genomic tessellations provide a canonical representation of repetitive and non-repetitive DNA and enable efficient sequence analyses while preserving homology.

**Software availability:** https://github.com/ekimb/tessellate.git

## 1 Introduction

A tessellation is a covering of a plane with a finite set of patterns, or *tiles*, derived from a single shape such that the tiles intersect only on their boundaries with no overlaps and no gaps. Historically, tessellations have been used for decorative effect in quilting and architecture. In Roman *opus tessellatum*, individual tesserae were assembled into decorative and figurative compositions based on an earlier Greek tradition of pebble mosaics [10]. M.C. Escher’s study of the interlocking motifs in The Alhambra inspired a series of pieces known as *Escher’s tilings* [15], in which repetition acts along one coordinate: translating by the length of a unit reproduces the same periodic template. Beginning the interval halfway through the unit changes its phase (or *geometry*), while retaining its cyclic *identity*.

DNA provides ample biological reason to make this distinction. Variation in short tandem repeats (STRs), which contain nearly-identical copies of a sequence unit, is associated with gene regulation [13] and disease [20]. Nowadays, third-generation sequencing technologies, such as PacBio HiFi and Oxford Nanopore (ONT) provide longer and more accurate reads that span and resolve periodic regions [7]. However, high overall read accuracy does not guarantee that sequencing errors are distributed uniformly. Errors in PacBio HiFi reads demonstrate systematic bias toward homopolymer length-errors and repeat-count errors in microsatellites, often supported by multiple reads [23]. Homopolymers and STRs are frequent sources of errors also in ONT reads, with longer array lengths underestimated systematically [17]; these errors are remedied by analysis of the duration for which the sequenced strand occupies the pore (dwell time) [28].

In order to mitigate the effect of error within periodicity, one fundamental case of this distinction has been well studied under *homopolymer compression* (HPC), which replaces each maximal run of identical nucleotides, for example, `AAAAA…A` with a single representative base `A`. Prior work demonstrated that separating the identity of the repeated nucleotide from its multiplicity improves sequence analysis tasks: `minimap2`, the state-of-the-art long-read mapper, uses homopolymer-compressed seeds [18]. HiCanu [23] collapses each homopolymer before read correction and overlap filtering to avoid penalizing run-length differences. Shasta [29] stores nucleotide identities and run lengths separately, constructs its assembly graph using markers from the compressed sequence, and estimates run lengths during the consensus step. Mapping-friendly sequence reductions [2] generalize HPC through a systematically-selected set of local transformations applied to both reads and reference sequences to improve mapping accuracy. Verkko constructs and resolves its assembly graph in homopolymer-compressed space before restoring homopolymers during final consensus [27].

We propose the paradigm of a *genomic tessellation* as a canonical representation of DNA that separates the atomic component (hereinafter referred to as the *identity*) and the way the component presents itself (the *geometry*) of any DNA sequence S. The identity of S is a canonical representative of all cyclic rotations of S, and the geometry of S describes how the identity appears throughout S, or equivalently, the operations that transform the identity back to S. For example, three exact realizations of identity `AC` are the sequences `ACACAC`, ACACACAC, `CACACA`, differing only in how `AC` is rotated and copied across each sequence. We show that we can construct tessellations in time linear to the size of the input such that sequences that are tiled with the same identity are separated by a bounded number of geometric differences, and that any sequence tiles identically to any substring thereof within a bounded neighborhood of the substring endpoints. We propose a compressed full-text index, the tessellated Burrows–Wheeler transform (tBWT), which enables exact matching queries on a reference tessellation, with auxiliary access to geometric differences. We demonstrate that the dichotomy between identity and geometry, captured by tessellations, and separating geometric differences from tile identity allows for efficient repeat-agnostic matching across homologous sequences with applications to read mapping, assembly polishing, and repeat resolution.

## 2 Methods

In this section, we formalize genomic tessellations as a representation of DNA that separates identity from geometry. An overview of the paradigm is provided in Figure 1. Supplementary text and proofs are provided in Appendix A.

**Figure 1:**
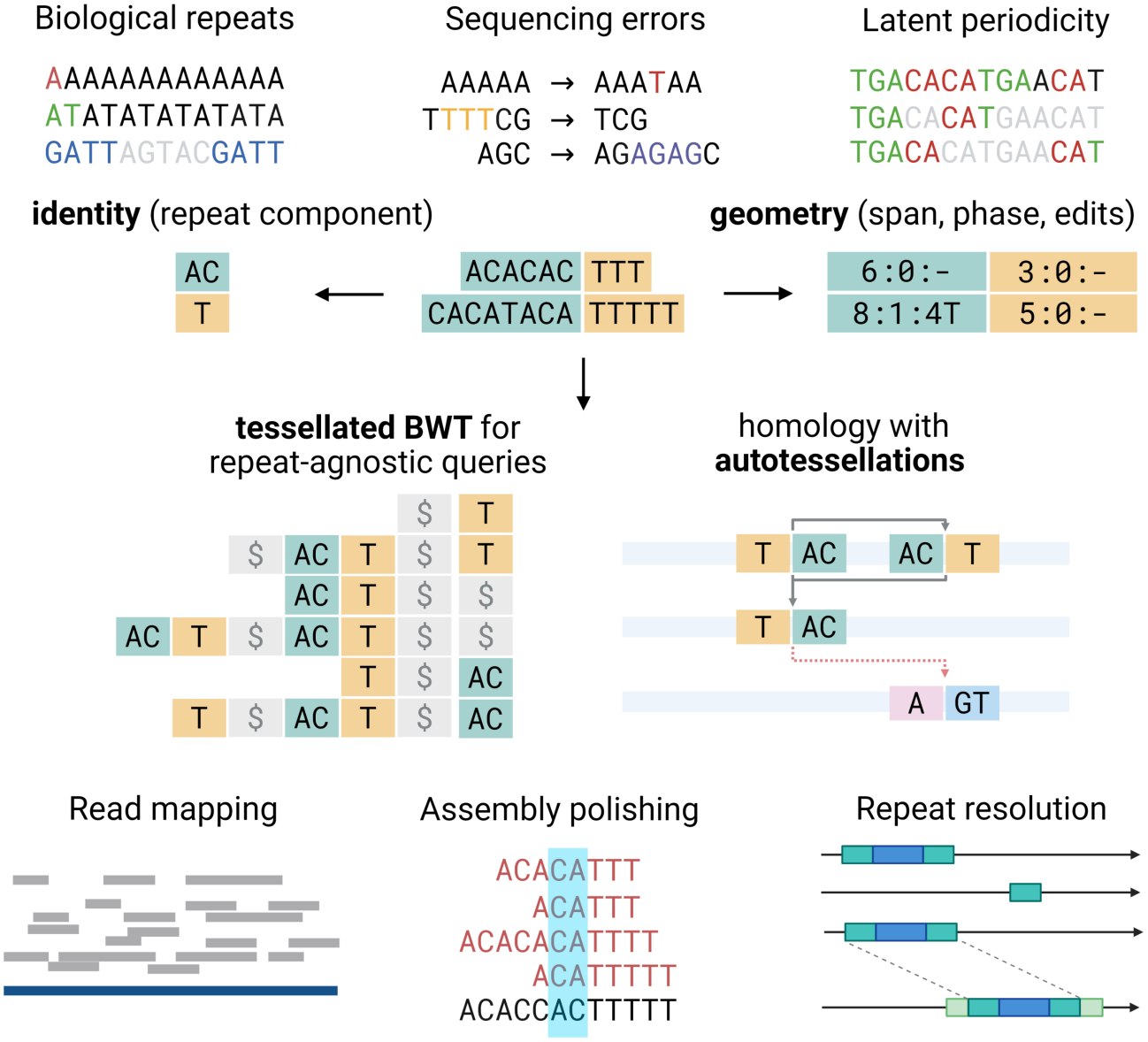
An overview of genomic tessellations and the tBWT. Periodicity is pervasive in genomic sequences in the form of known homopolymer or short tandem repeat (STR) arrays, or as frequent array-length and homopolymer errors in sequencing data, towards which third-generation sequencing technologies are systematically biased (top). A genomic tessellation (middle) is a lossless factorization of any DNA sequence into tiles each with a separate *identity* (middle, right) of a recurring component, and *geometry* of differences in how the component is presented. A tessellated Burrows–Wheeler transform (tBWT, middle, left) can be built on tile identities, with geometry stored and retrieved lazily. An *autotessellation* of a set of sequences is a set of correspondences built from the set of maximally-shared tile intervals on the tBWT that can be used, along with known geometric differences, to accelerate and improve read mapping, correcting misassemblies, and resolving repeats (bottom).

## 2 Methods

In this section, we formalize genomic tessellations as a representation of DNA that separates identity from geometry. An overview of the paradigm is provided in Figure 1. Supplementary text and proofs are provided in Appendix A.

### Definition 1

(alphabet, word). Let Σ_DNA_ = `A`, `C`, `G`, `T` be the DNA alphabet of nucleotides with order `A` *<* `C` *<* `G` *<* `T`. A *word* is a finite sequence of symbols from Σ we write Σ ^*^ to denote all such words, including the empty word ω, and Σ^+^ = Σ ^*^ *\ {ε}*. A sequence *S ∈* Σ^*n*^ has length |*S*| = *n* and symbols *S*_0_,. .., *S*_*n−*1_. We use zero-based, half-open intervals *S*[*a* : *b*) = *S*_*a*_*S*_*a*+1_ *S*_*b−*1_ for 0≤ *a ≤b ≤n*, with *S*[*a* : *a*) = ε. The word *xy* denotes concatenation of words *x* and *y*, and *w*^*c*^ denotes *c* concatenated copies of *w* for an integer *c ≥*0, with *w*^0^ = ε.

### Definition 2

(Reverse complementation). We denote complementation of a base *b* by 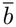, with `Ā = T` 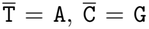 and 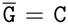. The *reverse complement S*^*†*^ of *S* reverses its order and complements each base: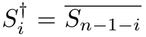, and ε^*†*^ = ε. For any word *w*, we say that *w* is *reverse-complement palindromic* if and only if *w* = *w*^*†*^ (for example, `ACGT`). This property is distinct from being a palindrome under reversal.

### Definition 3

(rotation, conjugacy class). For a word *w* = ε of length *p* and *j ∈* Z, we denote its *rotation* by *j* positions as

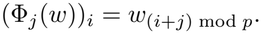

By convention, rotation amounts are interpreted modulo *p*, and positive rotations are left rotations. A word *w* is a *conjugate* of a word *v* if there exist *x, y ∈ Σ* ^*\**^ such that *w* = *xy* and *v* = *yx*. This is an equivalence relation whose *conjugacy class* is

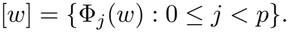

We identify each conjugacy class with its reverse-complement class to obtain the *strand-independent class* [*w*]_*±*_ = [*w*] ∪ [*w*^*†*^]. For example, [`ACG`] = *{*`ACG`, `CGA`, `GAC`*}*, and [`ACG`]_*±*_ = *{*`ACG`, `CGA`, `GAC`, `CGT`, `GTC`, `TCG`*}*.

### Definition 4

(period, primitivity, root). An integer 1 ≤ *q ≤* |*x*| is a *period* of a nonempty finite word *x* if *x*_*i*_ = *x*_*i*+*q*_ whenever 0 ≤ *i <* |*x*| *− q*. A nonempty word *w* is *primitive* if *w v*^*c*^ for every shorter nonempty word *v* and integer *c≥* 2. The *primitive root* of *w* is the shortest nonempty word *v* such that *w* = *v*^*c*^ for some integer *c ≥*1; for example, `ACACAC` has primitive root `AC`. A primitive word has exactly *w* distinct rotations.

### Definition 5

(identity, phase, dictionary). For fixed *P ≥*1 and a primitive word *w* of length *p ≥P*, let *a* = min[*w*] and *b* = min[*w*^*†*^] under lexicographic ordering. The *identity* ε(*w*) of *w* is defined by

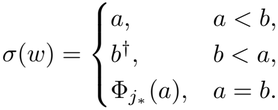

The last case chooses a reverse-complement-palindromic rotation with the smallest index, i.e., *j*_*\**_ is the minimum *j ∈ {*0,.. .,*p −* 1*}* satisfying Φ_*j*_(*a*)^*†*^ = Φ_*j*_(*a*). The *phase* of a primitive word *w* relative to its identity σ (*w*) is ϕ (*w*) ∈ *{*0,…, |*w*| *−* 1*}* satisfying *w* = Φ ϕ _(*w*)_(σ (*w*)). We define the *dictionary*

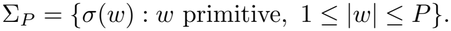

as the set of all identities for words of length ≤*P*. We prove in Appendix A.1 that there exists a unique identity and phase for each primitive *w* (Lemma 1). For example, when *P ≥*3, `ACG` and `CGA` have identity `ACG` and phases 0 and 1; their opposite orientation has identity `CGT` = `ACG`^*†*^. An identity need not be the lexicographically smallest rotation of its own class: the representatives are chosen across strands, and distinct orientations share an identity exactly when their rotation class is invariant under reverse complementation.

### Definition 6

(template). For identity σ ∈ Σ_*P*_ of length *p*, phase 0 *≤ φ < p*, and span *s ≥* 0, the *template*

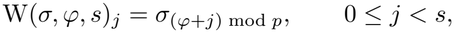

is the *s*-prefix of the infinite periodic expansion of *σ* starting at phase *φ* . In other words, W rotates the word *σ* by *φ*and repeats the resulting word until it reaches length *s*; for example, W(`AC`, 1, 5) = `CACAC`.

### Definition 7

(tile). A *tile t* is a representation of any DNA sequence *S ∈ Σ*^*\**^ of length *n* as an identity *σ* _*t*_ *∈ Σ*_*P*_ and *geometry γ*_*t*_ = (*s* _*t*_, *φ* _*t*_, *χ*_*t*_), where *s*_*t*_ = *n* is the observed span and *χ*_*t*_ is a sequence of *exceptions* (ordered edit operations). Applying *χ*_*t*_ to the template W(*σ*_*t*_, *φ* _*t*_, *ℓ*_*t*_) yields exactly *S*, where *ℓ*_*t*_ = *s*_*t*_ +*D*_*t*_ *− I*_*t*_ is the template length and *I*_*t*_, *D*_*t*_ count the bases inserted and deleted by the script. Below is an example set of sequences along with each of their tile identity and geometry for *P* ≥ 4 (exception positions are 0-based).

| S | σ<sub>t</sub> | s<sub>t</sub> | φ<sub>t</sub> | 𝒳<sub>t</sub> |
|---|---|---|---|---|
| ACACAC | AC | 6 | 0 | – |
| CACACA | AC | 6 | 1 | – |
| ACATACAC | AC | 8 | 0 | T : X : 3 |
| ACGT | ACGT | 4 | 0 | – |

Note that short, non-periodic sequences (such as `ACGT` above, whose tile identity is itself) and long, repetitive arrays both admit a tile representation, and that *S* can be recovered directly from t by generating the template W(*t*) = W(σ_*t*_, φ_*t*_, *l*_*t*_) and applying the exceptions *X*_*t*_. We denote the output of this procedure as ∥*t*∥ . For a prescribed *k* ≥1, we show that our construction in Subsection 2.1 deterministically produces tiles whose exception scripts have elementary cost at most k (Proposition 2 in Appendix A.1). We show in Appendix A.1 that any two tiles sharing an identity are related by at most 2k + 2 geometric adjustments (at most 2k elementary edits, one phase change, and one template-length change, Theorem 3). In Appendix A.3, we prove that reverse complementation is an involution on tiles, satisfies ∥t^*†*^∥ = ∥t∥^*†*^, and can be computed in *O*(*k*) operations for fixed maximum unit length *P* (Theorem 4).

### Definition 8

(tessellation). Let *S* ∈ Σ ^*^ be any sequence of length *n* and *A* = *a*_0_, *a*_1_,…, *a*_*m*_ be any strictly increasing sequence (a partition) over the interval [0, *n*]. For *k* ≥ 0, a *tessellation* of S is a sequence *τ* _*k*_(*S*) = *t*_0_*t*_1_ ·· · *t*_*m*−1_ such that

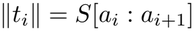

In other words, a tessellation *τ* _*k*_(*S*) is a factorization of any sequence S into tiles such that every position on S belongs to exactly one tile on *τ* _*k*_(*S*).

### 2.1 Linear-time tessellation of DNA sequences

A DNA sequence may admit periodic explanations with different identities. For example, `ACACAC` can be represented by identity `AC` without exceptions, or by identity `AAC` at phase 1, deleting positions 3 and 6 from ACAACAAC. For a prescribed error bound *k* ≥0, we seek a selection scheme for periodic explanations so that all selected explanations of the same sequence share one identity, and each requires at most *k* edits to recover the sequence from its periodic expansion. Locality-sensitive bucketing (LSB) provides a related guarantee based on pairwise edit distance: sequences within distance *d*_1_ share at least one bucket whereas sequences at distance at least *d*_2_ have disjoint bucket sets [5]. Our construction, instead, allows sequences with arbitrarily large pairwise edit distance to share an identity when their differences arise from length, or phase. For example, `ACAC` and ACACACACAC share identity `AC` despite an edit distance of 6. We prove in Appendix A.1 that all accepted periodic explanations of the same observed sequence agree on its identity, while their phase, span, and exceptions remain in geometry (Theorem 2); short exact fallback tiles retain exact non-periodic tile representations.

Given a fixed maximum unit length *P* ≥4 and prescribed error bound 0 ≤ *k* ≤ *K*, Algorithm 1 tessellates a sequence *S* of length *n* using a precomputed dictionary Σ_*P*_ and budget table *β*_*K*_. We start by scanning *S* for its maximal exact periodic runs of period *p* ≤ *P* (lines 4–10). Each list *C*_*p*_ for each period *p* ≤ *P* contains the maximum p-periodic intervals on S. Generate (*S, C*) (line 12; Algorithm 5 in Appendix A.2) merges and deduplicates the intervals in *C*, resolves run overlaps using a fixed ranking of periodic identities, and returns the set *Q* of *proposals q* = (*a*; *t*) (holding starting index a and tile t), gap intervals A = (z_0_,. .., z_|A|*−*1_), and boundaries b = (*b*_0_,. .., *b*_|A|_). Next, we propose extensions of periodic tiles into neighboring pieces and joins between nearby tiles sharing an identity (lines 14–27). A proposal *q* = (*a*; *t*) replaces consecutive raw pieces starting at a by tile t. We order proposals by a context-independent priority key of template length, edit cost, and a fixed strand-symmetric hash of the identity, retaining only those strictly better than every overlapping proposal in the original set. Finally, we emit the selected proposals and all untiled exact pieces (lines 29–35).

#### Algorithm 1

Tessellating a DNA sequence

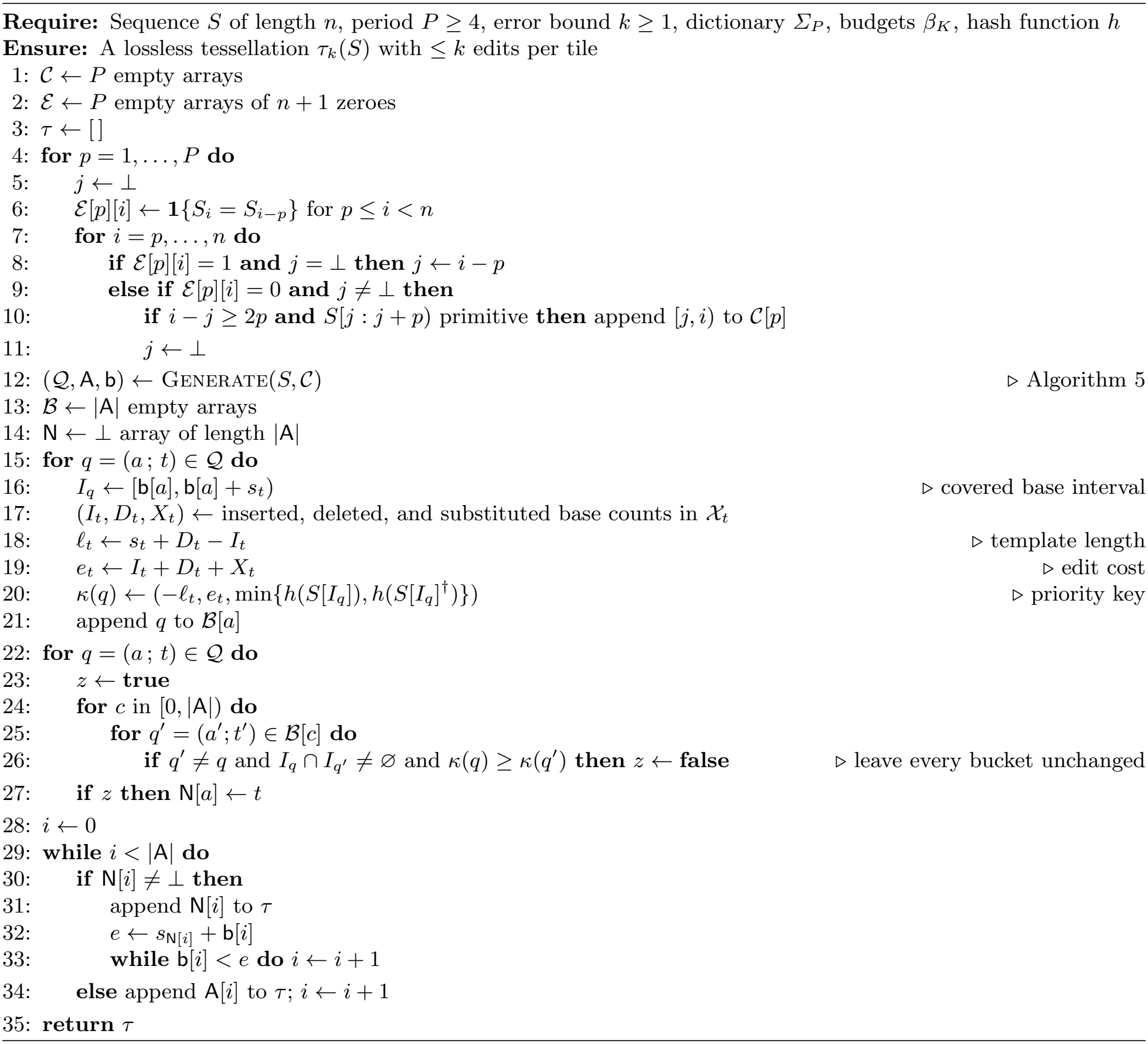

The resulting tessellation *τ*_*k*_(*S*) = (*t*_0_,. .., *t*_*m−*1_) satisfies ∥*t*_0_ ∥…∥*t*_*m−*1_∥ = *S*, and every tile has elementary edit cost at most *k* (Proposition 2). Construction takes *O*(*n*) time and space for fixed P and supported cap K (Proposition 3). Both the representation and the construction is invariant under reverse complementation: mirroring a tile satisfies (*t*^*†*^)^*†*^ = *t* and ∥*t*∥^*†*^ = ∥*t∥* ^*†*^ (Theorem 4), and independently tessellating the reverse strand gives *τ*_*k*_(*S*^*†*^) = (*τ*_*k*_(*S*))^*†*^ (Theorem 5). Explicit bounds and proofs are provided in Appendices A.2 and A.3.

### 2.2 The tessellated Burrows–Wheeler transform (tBWT)

By separating tile identity from tile geometry, tessellations permit exact matching across variations in repeat extent, rotations, and edit operations. We demonstrate that a compressed full-text index, based on the well-studied Burrows–Wheeler transform (BWT) [3, 11], can be constructed to count and locate the occurrences of tile identities *q* = *σ*_0_ .. . *σ*_*m*_, with geometry retrieved afterwards. Given *k* ≤ *K* and *P* ≥ 4, the *tessellated Burrows–Wheeler transform* (tBWT) indexes text *T* through a tessellation T = *τ*_*k*_(*T*) whose symbols are unit identities *σ* ∈ Σ_*P*_ . Queries on the tBWT are strictly on tile identity: for example, `locate`(`AT` *·* `G`) returns the positions of the occurrences of tiles *t*_AT_*t*_G_ on T with identity `AT` followed by identity `G` (for example, `ATG`, `ATATG`, `ATATGG`, `ATGGGG`). Note that this is *not* equivalent to finding all occurrences of form `AT`^*x*^`G`^*y*^ on *T*, since there is no guarantee that every substring of *T* with such form will exactly have the tiles *t*_AT_*t*_G_; instead, we establish in Subsection 2.1 the inverse relation that if there is a tile identity match between *τ*_*k*_(*q*) and T, the corresponding substrings on *q* and *T* share the same periodic explanation with at most *O*(*k*) modifications.

#### Definition 9

(tessellated text, tBWT)

For a sequence *S*, let U(*τ*_*S*_) = *σ*_0_ …*σ*_*m*_ be a function that generates an identity sequence from a tessellation of *S* into *m* tiles. For *M* input sequences *S*_0_,…, *S*_*M™*1_ and sentinel $, let 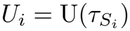. The *tessellated text* is

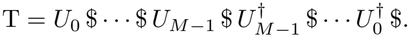

Analogously to its nucleotide counterpart, the tessellated BWT is the sequence

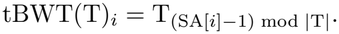

where SA is the suffix array over the tessellated text T. We store tBWT(T) in a wavelet tree with binary code depth at most *D*, supporting *O*(1) `access` and `rank` queries at each node. Since the unit alphabet is finite for fixed *P, D* = *O*_*P*_ (1). The following theorem immediately follows from [11] once tBWT(T) is equipped with an array of cumulative counts and a `rank` data structure:

#### Theorem 1

(count, locate)

*For a query of m tile identities, the tBWT counts its occurrences in O*(*mD*) = *O*_*P*_ (*m) time, and locates h such occurrences in O*(*D*(*m + hr)*) = *O*_*P*_ (*m + hr) time using O*(1) *auxiliary words*.

*W*e store tile geometry separately in blocks of *rc*onsecutive tiles aligned with the text positions sampled for `locate`. Each block encodes the span excess *δ*_*t*_ = *s*_*t*_ ™ |*σ*_*t*_| ≥ 0 of each tile *t* using an Elias gamma code [8] for *δ*_*t*_ + 1, its phase *φ*_*t*_ using a bitmap marking nonzero values followed by their packed values, and _*t*_ *X* using gap-encoded coordinates and packed operation codes. Two Elias–Fano directories [8, 9] store the blocks’ base coordinates and byte offsets, permitting access to individual blocks. A tile at a located offset *j t*hen lies in block ⌊*j/r*⌋, and its base coordinate is the start position of its block plus the preceding spans *s*_*t*_ = |*σ*_*t*_ |+ *δ*_*t*_ within the block. Reverse-strand hits can equivalently be recovered using mirrored offsets and the tile reverse-complement operation of Theorem 4.

A tessellated text built from a set of input sequences may contain tile identities that recur within the same sequence or in multiple sequences. Matching the tessellated text against itself through tBWT queries makes this correspondence explicit while retaining tile geometry for each occurrence. We call this operation an *autotessellation*, which we obtain by building the longest-common-prefix (LCP) array for tessellated text T, and recording the exact matches that admit a correspondence. Let *R =* (*S*_0_,…, *S*_*∥™*1_) be a set of sequences with tessellated text T of length *N* built from identity sequences *U* _*i*_ = U(*τ*_*i*_), *m*_*i*_ = |*U* _*i*_|, and *F*_*r*_ the suffix of row *r* of tBWT(T). The *longest common prefix* (LCP) array is

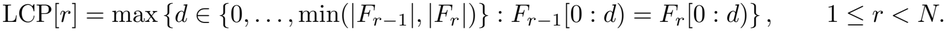

The LCP array can be constructed directly from tBWT(T) using a BWT-based algorithm [1] extended to string collections [26]. The construction traverses selected BWT intervals using backward extensions and obtains LCP values from the lengths of the strings represented by those intervals, which can be performed in *O* (*M*log |Σ_*P*_|) time with the same wavelet-tree representation as that of the tBWT. Alternatively, when the tessellated text and the complete suffix array are available during tBWT construction, the LCP array can be computed in *O*(*M*) time using the algorithm by Kasai *et al*. [16]. For the suffix beginning at tile *a* of *U* _*i*_, let *r*_*i*_(*a*) be its tBWT row. Its longest prefix occurring at another indexed placement has length

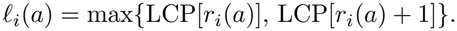

Since a suffix attains its longest common prefix with another suffix at a lexicographic neighbor, it suffices to examine the adjacent rows. Retaining the positive-length intervals [*a, a*+*l*_*i*_(*a*)) that are not strictly contained in another interval gives the supermaximal query intervals, which are recovered with `locate` queries on the tBWT. The *autotessellation* of the input collection *R* is

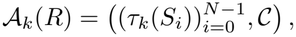

where *C* contains every correspondence (*i, a, j, b, d, λ*) satisfying maximal *a* with

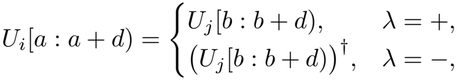

We demonstrate later in Subsection 3.3 that autotessellations capture homology among input sequences by locating all occurrences of maximal repeated identity intervals while retaining the geometry of each occurrence through the original tessellations.

## 3 Results

We used the HG002 Q100v1.1 benchmark [14] throughout our evaluations, for which large long-read sequencing data is readily available. We used `hifiasm` [6] as the long-read assembler using both ONT-only parameters (`--ont`) and HiFi with ultra-long ONT integration (`--ul`). Throughout all experiments, we used `minimap2` for comparing draft assemblies to the Q100 reference.

### 3.1 Tessellations outperform state-of-the-art long-read mappers in repeat-dense regions

Long-read mapping is known to be particularly challenging in repeat-dense regions, such as satellite arrays: highly-frequent seeds are discarded or down-weighted to keep chaining tractable, and the reads originating from an array are unmapped or have low mapping quality (mapQ). Tessellations mitigate these challenges in two ways: First, tiles absorb small differences that dominate long-read sequencing errors so that reads and their true loci share longer exact matches in tile identity than in nucleotides. Second, exact tile identity matches, remarkably, allow seeding without an occurrence threshold: every occurrence of each match can be efficiently located and ranked based on the tiles at which the read tile differs from the reference autotessellation.

We evaluated read mapping on a synthetic reference from the HG002 Q100v1.0 maternal X chromosome (`chrX MATERNAL`, 154.34Mbp), and expanded two of its tandem arrays, the centromeric *α*-satellite array DXZ1 and macrosatellite DXZ4, by mutating each repeat unit copy independently with a constant divergence rate using `simulate tandem repeat.py` from `centroFlye`[4]. We simulated 500 additional copies of the ∼ 2kbp DXZ1 repeat unit each at 0.5% divergence and inserted the simulated units into the native array on chrX (57.8 – 60.9Mbp) at the boundary nearest the array’s midpoint, expanding it to ∼ 4.1Mbp. For DXZ4, we extracted a 2.97kbp repeat unit from the native array on chrX (114.22 — 114.44Mbp), and simulated and inserted 50 copies at 0.5% divergence at its midpoint. We simulated ONT reads from the array using `nanosim` [30], and with the R10.4.1 pore model (error rate 1.4%, median length 11kbp) at 20× depth, and PacBio HiFi reads with `pbsim2` [24] With ONT R10.4.1 reads (top), `tessellate`, `minimap2` and `Winnowmap2` misplace 260, 286 and 225 of the 6,150 reads from the DXZ1 region. tessellate assigns mapQ 60 to 5,164 DXZ1-region reads perfectly, against 3,216 and 66 for `Winnowmap2` and `minimap2`, respectively.

With PacBio HiFi reads (bottom), the arrays are more tractable for seed-chain-extend-based mappers. `Winnowmap2` and `tessellate` have comparable confidence in the DXZ1 array, with `Winnowmap2` mapping 4,628 reads at mapQ 60 against 4,611 for `tessellate`, and 361 for `minimap2`. On the DXZ4 array, `tessellate` gives mapQ 60 to 362 reads, against 238 for `Winnowmap2` and 28 for `minimap2`, with one wrong placement. However, long-read mapping with tessellations matches the other mappers at a fraction of their cost: `tessellate` maps the 20x ONT read set 57x and 60x faster than `minimap2` and `Winnowmap2`, respectively, using ∼ 5x less memory. On the HiFi read set, `tessellate` is 27x and 83x faster than `minimap2` and `Winnowmap2`, respectively, while using 4x–6x less memory.

**Table 1:** Mapping statistics of tessellated exact matches, `minimap2`, and `Winnowmap2` on the maternal X chromosome of HG002 with synthetic expansions of the DXZ1 and DXZ4 arrays. All tools were run a AMD EPYC 9575F using 64 threads without nucleotide-level alignment; reported overlaps were evaluated with `paftools mapeval`. For Winnowmap2, the time for reference *k*-mer counting (`meryl count`) was not included.

| Technology | Method | Region | Q60 mapped | Q60 wrong | mapped | wrong | unmapped | Runtime (s) | Memory (GB) |
|---|---|---|---|---|---|---|---|---|---|
| ONT R10.4 20× | minimap2 | DXZ1 | 66 | 0 | 6125 | 286 | 25 | 982.6 | 34.1 |
|  |  | DXZ4 | 39 | 0 | 623 | 36 | 0 |  |  |
|  |  | all | 180310 | 0 | 188924 | 706 | 41 |  |  |
|  | winnowmap2 | DXZ1 | 3216 | 0 | 6149 | 225 | 0 | 1029.1 | 48.3 |
|  |  | DXZ4 | 258 | 0 | 623 | 51 | 1 |  |  |
|  |  | all | **183737** | 1 | 188948 | 697 | 17 |  |  |
|  | tessellate | DXZ1 | **5164** | 0 | **6150** | 260 | 0 | **17.1** | **8.6** |
|  |  | DXZ4 | **460** | 0 | 623 | 42 | 0 |  |  |
|  |  | all | 177230 | 2 | **188954** | **692** | 11 |  |  |
| PacBio HiFi 20× | minimap2 | DXZ1 | 361 | 0 | 4765 | 2 | 0 | 296.9 | 25.8 |
|  |  | DXZ4 | 28 | 0 | 422 | 0 | 0 |  |  |
|  |  | all | 169088 | 0 | 176197 | **156** | 0 |  |  |
|  | winnowmap2 | DXZ1 | **4628** | 0 | 4765 | 1 | 0 | 912.3 | 40.6 |
|  |  | DXZ4 | 238 | 0 | 422 | 0 | 0 |  |  |
|  |  | all | **174664** | 0 | 176197 | 160 | 0 |  |  |
|  | tessellate | DXZ1 | 4611 | 0 | 4765 | 1 | 0 | **11.2** | **7.4** |
|  |  | DXZ4 | **362** | 1 | 422 | 3 | 0 |  |  |
|  |  | all | 174040 | 1 | 176197 | 181 | 0 |  |  |

### 3.2 Tessellations efficiently capture and correct small assembly errors

Small errors in draft assemblies frequently occur around repetitive regions [22], and are typically corrected by specialized variant callers and polishers that require a full alignment of HiFi or ONT reads to the assembly. In this section, we demonstrate that an autotessellation of the assembly, together with a tessellation of the reads, serves as a direct substitute for this task.

`tessellate polish` replaces read-to-assembly alignment with exact matches in tile space. We tessellate both the assembly and reads and build an autotessellation (described in Subsection 2.2) of the reference. Each read is then placed by locating its super-maximal exact matches (SMEMs [19]) of at least k tiles in the index (k = 12 by default). Finding SMEMs between the read tessellations and the reference autotessellation reports, per assembly tile, the reads that contain a SMEM at that interval, which, by definition, differ from the assembly only in geometry. Tiles where the fraction of agreeing reads below the median agreement across the autotessellation are flagged as errors, and neighbouring candidates are merged into windows bounded by tiles supported by the reads. Finally, the window is replaced by geometry the reads most frequently support. `tessellate` also exposes copy-number errors using the repeat table by removing a copy whose sequence, junctions included, is not supported by any read.

We autotessellated two draft assemblies of the maternal X chromosome of HG002: a `hifiasm` assembly built only from ONT reads carrying ∼14k small errors provided by [14], and a hybrid `hifiasm` PacBio HiFi assembly with ultra-long ONT (UL) integration carrying fewer than 700. For the ONT-only assembly, the evaluation against Q100 [14] reports 44.6kb of extra DXZ4 sequence as a structural discrepancy. For the hybrid assembly, our alignment to the Q100 reference shows insertions of 4.1kb within DXZ1 and 44.6kb within DXZ4. We polished each assembly with `tessellate`, `DeepVariant` [25] and `DeepPolisher` [21], using the same HiFi read set of ∼54 x depth. Table 2 shows the evaluation of each polished assembly against the HG002 Q100 v1.1 reference in terms of percent identity (% to Q100), base-level and event-level QV, *k*-mer QV calculated with `yak`^1^, and runtime and memory.

**Table 2:** Polishing long-read assemblies, evaluated on two `hifiasm` assemblies of the maternal X chromosome of HG002 against the Q100 reference. The “Differences” column denotes the number of SNVs, insertions and deletions in the alignments of each assembly to the Q100 reference (`minimap2 -x asm5`). “Fixed” and “new” columns are difference counts relative to the unpolished assembly’s differences. “Identity” denotes the percentage of matching bases over alignment columns. *k*-mer QV is calculated using `yak -k 31`. Event QV is −10 log_10_(differences / columns), and base QV is −10 log_10_(edit distance / columns). All methods used the same HiFi reads. Best values for each assembly are in bold.

| Assembly | Method | Differences | Fixed / new | Identity (%) | *k*-mer QV | Event QV | Base QV | Runtime (s) | Memory (GB) |
|---|---|---|---|---|---|---|---|---|---|
| ONT-only | Unpolished | 14,103 | – | 99.959 | 45.39 | 40.40 | 33.85 | – | – |
|  | DeepPolisher | 4,012 | 11,174 / 1,105 | 99.968 | 50.73 | 45.86 | 34.94 | 1983.1 | 29.8 |
|  | DeepVariant | **398** | 13,710 / 29 | 99.971 | **62.42** | **55.89** | 35.32 | 311.3 | 29.2 |
|  | tessellate polish | 490 | **13,722** / 117 | **99.999** | 60.64 | 54.99 | **52.01** | **55.7** | **10.0** |
| Hybrid | Unpolished | 702 | – | 99.968 | 57.91 | 53.42 | 34.90 | – | – |
|  | DeepPolisher | 2,260 | 198 / 1,756 | 99.966 | 52.41 | 48.35 | 34.75 | 2035.9 | 29.9 |
|  | DeepVariant | **518** | **216** / 35 | 99.968 | **59.05** | **54.74** | 34.92 | 279.7 | 29.0 |
|  | tessellate polish | 593 | 210 / 100 | **99.999** | 57.73 | 54.15 | **51.11** | **43.0** | **9.3** |

On the ONT-only assembly, `tessellate` fixes 13,722 of the 14,074 original errors, as many as `DeepVariant` (13,710) and far more than `DeepPolisher` (11,174), which, trained on HiFi-based diploid assemblies, adapts poorly to haploid assemblies. The base-level QV improvement of `tessellate` originates from its correction of expanded satellite arrays. Estimating the copy number of each correspondence interval on the autotes-sellation from HiFi read support removes the extra DXZ4 sequence in both assemblies and the extra DXZ1 sequence in the hybrid. This raises identity to 99.999% and QV to ∼52. These corrections are not detected in *k*-mer QV values, since an extra array copy does not change the number of *k*-mers absent from the truth. In addition, `tessellate polish` runs under a minute using 10 GB (without a GPU), compared to ∼5 minutes with `DeepVariant` and about 34 minutes with a GPU with `DeepPolisher`, using ∼30 GB of maximum memory during read alignment.

### 3.3 Tessellations capture higher-order homology in fragmented assemblies

Figure 2 shows the generation of a single complete assembled fragment of the HG002 mitochondrial chromosome from an autotessellation of the `hifiasm` ONT-only assembly. We found exact tile-identity matches from a 20x set of HiFi reads and collected *links* from exact matches that span multiple fragments that share correspondence in the autotessellation: the largest combined link weight from the reads are within 16,550–16,600bp between the contigs `h2tg000157l` and `h2tg000095l`. Retrieving geometric differences within exact matches at this interval revealed that only 1 of 2,749 reads matched the assembly sequence GAGAGAGA, with 2,735 supporting `GAGAGA`. We applied this two-base contraction, which produced a 16,570-bp sequence matching the complete Q100 mitochondrial reference exactly after accounting for strand and circular origin, increasing gap-inclusive identity from 99.98% to 100%.

**Figure 2:**
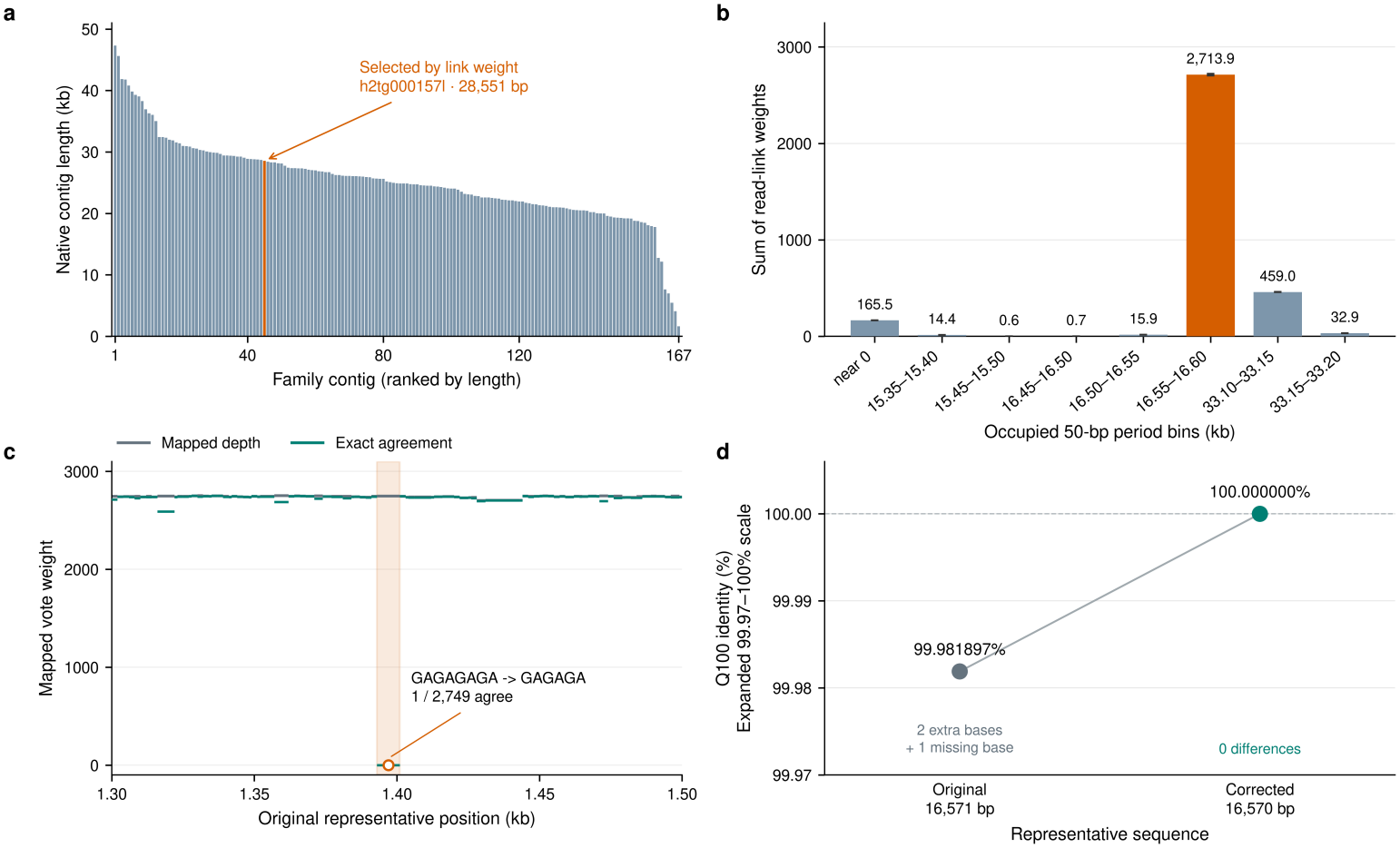
Generating a single consensus from a fragmented assembly of the human mitochondrial genome. Among 167 contigs ranked by length, the representative (orange) is selected by the strongest read-supported connection between contig ends within the most strongly supported repeat-length range (a). Estimated lengths of a repeating mitochondrial sequence unit are grouped into 50-bp ranges, and their summed read-link weights identify 16,550–16,600 bp as the most strongly supported range (b). Mapped vote weight (gray) and exact-agreement weight (green) reveal a discrepancy at the highlighted GA repeat, where only 1 of 2,749 reads matches the assembly and 2,735 support shortening GAGAGAGA to `GAGAGA` (c). Removing one `GA` repeat improves identity to the complete 16,570bp Q100 mitochondrial reference from 99.98% to 100% (d).

## 4 Discussion

Genomic tessellations are a lossless representation of both repetitive and non-repetitive DNA, and separate sequence identity from the geometry through which it is realized. This distinction allows sequence comparison to account for shared periodic structure while retaining repeat lengths, phases, and exceptions for downstream analysis. Autotessellation organizes sequence homology around shared tile identities while retaining geometric differences, which allows related intervals to remain recognizable through repeat-length changes and provides a common coordinate system for a correspondence family. The example in Section 3.3 illustrates how this organization can guide representative selection and correction within a family of redundant assembled fragments. Reliable interpretation still requires distinguishing assembly errors from genuine variation among copies, particularly when repetitive sequence permits multiple read placements.

There are many promising future directions where tessellations can mitigate challenges. Combining tile identity matches on the tBWT with geometry-aware sequence *alignment* may help better assess variation under sequencing errors. Furthermore, incorporating platform-specific error models, extending the unit dictionary to longer periods, and developing hierarchical autotessellations that capture recurring arrangements of tiles are natural extensions of this work. An open question is whether substitutions and indels can be shown to affect only a bounded number of tiles, with bounds expressed in terms of repeat period, allowed divergence, and ambiguity among competing decompositions. Moreover, determining the minimal sufficient conditions for recovering copy-number relationships and selecting representatives in terms of unique sequence content, length, and coverage remains open.

## Supporting information

Supplementary Text and Proofs

## Footnotes

1 https://github.com/lh3/yak

