## Supplementary Text and Proofs for "Genomic tessellations"

### A Supplementary text and proofs

#### A.1 Identity and error-tolerance

**Lemma 1 (identity is well-defined).** *For every primitive word $w$ of length $p \le P$, there exists unique $\sigma(w) \in [w]$ and unique $\phi(w) \in \{0, \ldots, p-1\}$ such that, for every $j \in \mathbb{Z}$,*

$$w = \Phi_{\phi(w)}(\sigma(w)), \qquad \sigma(\Phi_j(w)) = \sigma(w), \qquad \sigma(w^\dagger) = \sigma(w)^\dagger.$$

*Proof.* When $a \ne b$, Definition 5 is constant on $[w]$. When $a = b$, there exists some rotation $h$ such that $a^\dagger = \Phi_h(a)$. Such a rotation pairs positions $s \mapsto (p - 1 + h - s) \pmod p$, and since no base equals its complement, the position map has no fixed point. Hence, $p$ is even, and we can choose $j$ satisfying $2j \equiv h \pmod p$, which yields

$$(\Phi_j(a))^\dagger = \Phi_{-j}(a^\dagger) = \Phi_{h-j}(a) = \Phi_j(a).$$

Thus, a reverse-complement-palindromic rotation $j_*$ exists, is a unique representative determined by the conjugacy class, and unchanged by reverse complementation. Finally, primitivity implies that the $p$ rotations of $w$ are distinct; therefore, every other word $v \in [w]$ has unique $\phi(v)$.

**Definition 10 (eligible templates).** For the dictionary $\Sigma_P$, the family of eligible periodic templates is

$$\mathcal{W}_P = \{\mathrm{W}(\sigma, \varphi, \ell) : \sigma \in \Sigma_P, \quad 0 \le \varphi < |\sigma|, \quad \ell \ge 2|\sigma|\}.$$

for template length $\ell$. Eligibility requires at least two copies, but permits a partial final copy; for template $W = \mathrm{W}(\sigma, \varphi, \ell)$, the identity $\mathrm{I}(W) = \sigma$ is well-defined by Lemma 2.

**Definition 11 (separation, witness, certification).** For a prescribed error bound $k \ge 0$ and $W \in \mathcal{W}_P$, we define the *separation* $\Delta$ and *budget* $\beta_k$ as

$$\Delta(W) = \min_{\substack{V \in \mathcal{W}_P \\ \mathrm{I}(V) \ne \mathrm{I}(W)}} d(W, V), \qquad \beta_k(W) = \min\left\{k, \left\lfloor \frac{\Delta(W) - 1}{2} \right\rfloor\right\}.$$

A *witness* $(W, \mathcal{X})$ for an observed word $w$ is *certified* when $\mathcal{X}(W) = w$ and its elementary cost satisfies $e \le \beta_k(W)$. Certification ensures $e \le k$ and $2e < \Delta(W)$, even when the witness is not a minimum-distance alignment.

**Lemma 2 (template identity is well-defined).** *Every word $W \in \mathcal{W}_P$ has a unique identity $\iota(W)$, and $1 \le \Delta(W) < \infty$.*

*Proof.* Suppose the same word has two eligible primitive periods $p, q$. Its length is at least $2\max(p, q)$, so the Fine–Wilf periodicity theorem gives period $g = \gcd(p, q)$ [12]. The length-$p$ prefix consists of $p/g$ copies of its length-$g$ prefix, so primitivity forces $p = g$. Likewise $q = g$, and hence $p = q$; the phased units agree and have the same canonical identity. Different identities cannot give the same template word, so $\Delta(W) \ge 1$.

**Theorem 2 (unique certified identity).** *For a fixed observed word $x$, all certified witnesses have the same oriented identity. Moreover, if $(W, \mathcal{X})$ is certified with cost $e$, then every eligible template $V$ of another identity satisfies*

$$d(x, V) > e \ge d(x, W), \qquad e \le k.$$

*Proof.* Suppose two certified witnesses $W_1, W_2$ have different identities and costs $e_1, e_2$. Set $d = \mathrm{ed}(W_1, W_2)$. By separation, $2e_i < \Delta(W_i) \le d$ for $i = 1, 2$. The triangle inequality gives

$$d \le \mathrm{ed}(W_1, x) + \mathrm{ed}(x, W_2) \le e_1 + e_2 < d,$$

a contradiction. For any other-identity template $V$,

$$\mathrm{ed}(x, V) \ge \mathrm{ed}(W, V) - \mathrm{ed}(W, x) \ge \Delta(W) - e > e.$$

The witness itself gives $\mathrm{d}(x, W) \le e$, and $e \le \beta_k(W) \le k$ by certification.

**Definition 12 (Certified ball).** For $W \in \mathcal{W}_P$, define its certified ball by $B_k(W) = \{x \in \Sigma^* : \mathrm{d}(x, W) \leq \beta_k(W)\}$.

**Corollary 1 (Common-template stability).** *Balls belonging to different identities are disjoint. If $x, y \in B_k(W)$, every certified whole-interval description of either word has identity $\iota(W)$, and*

$$\mathrm{d}(x, y) \leq 2\beta_k(W) \leq 2k.$$

*Proof.* Minimum-distance scripts certify membership in each ball, so Theorem 2 implies disjointness across identities. For a common template, apply the triangle inequality through $W$.

**Lemma 3 (Separation grows with template length).** *If $W \in \mathcal{W}_P$ has primitive period $p \leq P$, then*

$$\Delta(W) \geq \left\lfloor \frac{|W|}{p + P - 1} \right\rfloor.$$

*Consequently, if $|W| \geq (2k+1)(p+P-1)$, then $\beta_k(W) = k$: every valid witness to $W$ costing at most $k$ edits is certified.*

*Proof.* Let $V$ be a different-identity rival of primitive period $q \leq P$, and put $B = p + P - 1$. An exact matching block of length at least $B$ would have both periods and length at least $p + q - \gcd(p, q)$. Fine–Wilf and primitivity would then force the same identity. Thus every exact matching block in an alignment of $W, V$ has length at most $B - 1$. An alignment of cost $d$ has at most $d + 1$ exact matching blocks. At most $d$ source-template bases are consumed by substitutions or deletions; insertions consume none. Hence

$$|W| \leq d + (d+1)(B-1) = (d+1)B - 1, \qquad d \geq \left\lfloor \frac{|W|}{B} \right\rfloor.$$

taking the minimum over rivals, at the stated threshold $\Delta(W) \geq 2k + 1$, which gives $\beta_k(W) = k$.

**Proposition 1 (Finite exact budget table).** *Fix a maximum supported error cap $K$. All radii $\beta_k(W)$ for $0 \leq k \leq K$ admit finite precomputation depending only on $P, K$.*

*Proof.* Put $L_* = (2K+1)(2P-1)$. For every eligible $W$ with $|W| \geq L_*$, Lemma 3 gives $\beta_k(W) = k$. Only eligible lengths below $L_*$ need explicit table entries. For such a template, compute the truncated separation $\delta(W) = \min\{\Delta(W), 2K+1\}$. A rival $V$ can lower this value only if

$$\big||V| - |W|\big| \leq 2K,$$

because edit distance is at least the length difference. There are finitely many primitive units, phases, and eligible rival lengths in this band. Global edit-distance dynamic programming computes their distances, capped at $2K+1$; the diagonal band $|i - j| \leq 2K$ suffices for every distance that can change the result. Store

$$b(W) = \left\lfloor \frac{\delta(W) - 1}{2} \right\rfloor.$$

Then $\beta_k(W) = \min\{k, b(W)\}$ for stored lengths and equals $k$ at or above the cutoff, and indexing provides $\mathcal{O}(1)$ lookup once the table is built.

**Proposition 2 (Deterministic lossless bounded-error output).** *Fix $P \geq 4$, $0 \leq k \leq K$. For every input $S$ of length $n$, the algorithm deterministically produces a tessellation $\tau_k(S)$. Its emitted tiles have positive spans totaling $n$, render $S$ exactly, and have elementary exception cost at most $k$. Every periodic replacement is certified by* (2).

*Proof.* The detailed construction is given in Appendix A.2, where the exact-run scans, overlap constraints, and gap cuts are determined by $S$. The gap-witness procedure $\mathit{\Gamma}$ uses a fixed orientation and a lexicographic minimum, or rejects the fragment, making emission in raw-piece order deterministic. The 4-nucleotide bound on gap-fragment lengths ensures that their roots belong to the dictionary, since $P \geq 4$ by definition. Every emitted tile is either an unchanged exact piece, whose script has cost 0, or a selected proposal $t$ that passed the acceptance test (2). In the latter case, with template $W_t$,

$$e_t \leq \beta_k(W_t) \leq k.$$

Selection and emission leave these scripts unchanged, so the bound holds for every output tile, including exact fallback tiles.

**Definition 13 (Geometric adjustment).** Fix an identity $\sigma$ of length $p$. A *geometric adjustment* is one of the following operations, each with unit cost:

1. an insertion, deletion, or substitution of one nucleotide;
2. a phase change from $\mathrm{W}(\sigma, \varphi, \ell)$ to $\mathrm{W}(\sigma, \varphi', \ell)$, with $\varphi, \varphi' \in \{0, \dots, p-1\}$;
3. a template-length change from $\mathrm{W}(\sigma, \varphi, \ell)$ to $\mathrm{W}(\sigma, \varphi, \ell')$, with $\ell, \ell' \geq 0$.

The last two operations act on exact templates and preserve $\sigma$. A phase change selects a different starting phase of the infinite periodic expansion, rather than rotating the finite template word. Each phase or length change counts as one adjustment regardless of its magnitude.

**Theorem 3 (Bounded geometric adjustments).** *Let $t, t'$ be tiles sharing the same oriented identity $\sigma$, with templates $W = \mathrm{W}(\sigma, \varphi, \ell)$ and $W' = \mathrm{W}(\sigma, \varphi', \ell')$. Suppose their exception scripts have elementary costs $e, e' \leq k$, counting every inserted, deleted, or substituted base. Then $\|t\|$ can be transformed into $\|t'\|$ using at most*

$$e + e' + \mathbf{1}\{\varphi \neq \varphi'\} + \mathbf{1}\{\ell \neq \ell'\} \leq 2k + 2$$

*geometric adjustments. In particular, at most $2k$ adjustments are elementary nucleotide edits, at most one changes phase, and at most one changes template length.*

*Proof.* Write $x = \|t\| = \mathcal{X}_t(W)$ and $x' = \|t'\| = \mathcal{X}_{t'}(W')$. Expanding any encoded edit runs in $\mathcal{X}_t$ into elementary operations and undoing these operations in reverse order transforms $x$ back into $W$ in $e$ edits. If $\varphi \neq \varphi'$, perform one phase adjustment to obtain $\mathrm{W}(\sigma, \varphi', \ell)$. If $\ell \neq \ell'$, perform one template-length adjustment to obtain $W'$. Finally, apply $\mathcal{X}_{t'}$ to obtain $x'$ in $e'$ elementary edits. Summing these costs gives the stated bound.

##### A.2 Construction details and analysis

Here, we describe the overlap resolution, gap completion, proposal generation, selection, and emission subroutines in Algorithm 1. We fix $P \geq 4$, a supported maximum error cap $K$, and $0 \leq k \leq K$. Canonical identities and phases are those of Definition 5. Before processing sequences, we precompute $B[\sigma, \varphi, \ell] = \beta_K(\mathrm{W}(\sigma, \varphi, \ell))$ for eligible template lengths $\ell < L_* = (2K+1)(2P-1)$, using Proposition 1. A query at error cap $k \leq K$ returns $\min\{k, B[\sigma, \varphi, \ell]\}$ below this $L_*$ and $k$ otherwise.

**Resolving overlaps.** Line 1 of Algorithm 5 applies Algorithm 2 to $S$ and the array $\mathcal{C}$ of per-period interval lists. We enumerate the lists in $\mathcal{C}$ and merge intervals by increasing start and, for equal starts, decreasing end, retaining each interval's period index $p$. Let $\chi$ be a rank on classes under rotation and reverse complementation so that

$$\chi(\mathit{\Phi}_j(w)) = \chi(w) = \chi(w^\dagger).$$

For every overlap of two original intervals, $[a, b) = [a_j, e_i)$ with $i < j$ and $a < b$, we accumulate the following constraints simultaneously:

$$\begin{array}{r|l} \chi(w_i) < \chi(w_j) & L_j \leftarrow \max(L_j, b), \\ \chi(w_i) > \chi(w_j) & R_i \leftarrow \min(R_i, a), \\ \chi(w_i) = \chi(w_j) & R_i \leftarrow \min(R_i, a + \lfloor (b-a)/2 \rfloor), \\ & L_j \leftarrow \max(L_j, b - \lfloor (b-a)/2 \rfloor). \end{array}$$

**Algorithm 2** RESOLVE: merge and filter run lists and resolve overlaps

**Require:** $S$, array $\mathcal{C}$ of interval lists; class rank $\chi$
**Ensure:** Records $(L_i, R_i, p_i, w_i')$ in start order

$\mathcal{D} \leftarrow$ merge $(\mathcal{C}[p])_{p=1}^{|\mathcal{C}|}$ by increasing start, then decreasing end
$\mathcal{F} \leftarrow [\,]$; $e_{\max} \leftarrow 0$
**for** each interval $[a, e)$ from source list $p$ in $\mathcal{D}$ **do**
    **if** $e > e_{\max}$ **then**
        $w \leftarrow S[a : a + p)$
        Append $(a, e, p, w)$ to $\mathcal{F}$; $e_{\max} \leftarrow e$
$c \leftarrow |\mathcal{F}|$; write $\mathcal{F} = ((a_i, e_i, p_i, w_i))_{i=0}^{c-1}$
**for** $i = 0, \ldots, c-1$ **do**
    $L_i \leftarrow a_i$; $R_i \leftarrow e_i$
**for** $i = 0, \ldots, c-1$ **do**
    **for** $j = i+1, \ldots, c-1$ **do**
        **if** $a_j \geq e_i$ **then break**
        $a \leftarrow a_j$; $b \leftarrow e_i$ ▷ use the original endpoints
        **if** $\chi(w_i) < \chi(w_j)$ **then**
            $L_j \leftarrow \max(L_j, b)$
        **else if** $\chi(w_i) > \chi(w_j)$ **then**
            $R_i \leftarrow \min(R_i, a)$
        **else**
            $h \leftarrow \lfloor (b-a)/2 \rfloor$
            $R_i \leftarrow \min(R_i, a+h)$; $L_j \leftarrow \max(L_j, b-h)$
$\mathcal{C}' \leftarrow [\,]$ ▷ discard intervals only after all comparisons
**for** $i = 0, \ldots, c-1$ **do**
    **if** $R_i - L_i \geq p_i$ **then**
        $w_i' \leftarrow \Phi_{L_i - a_i}(w_i)$ ▷ preserve the unit, advance its phase
        Append $(L_i, R_i, p_i, w_i')$ to $\mathcal{C}'$
**return** $\mathcal{C}'$

**Filling gaps.** After resolving overlaps, GENERATE completes the retained runs to an exact partition using Algorithm 3. Let $\rho$ be the function $\rho(\mathtt{A}) = 0, \rho(\mathtt{C}) = 1, \rho(\mathtt{G}) = 2, \rho(\mathtt{T}) = 3$. We partition each uncovered gap by its endpoints and by cuts isolating ambiguous bases, and cut

$$\begin{array}{rl} \text{at } i & \text{if } S_{i-1} = S_i, \\ \text{at } i & \text{if } \rho(S_{i-1}) < \rho(S_i) > \rho(S_{i+1}), \\ \text{at } i+1 & \text{if } \rho(S_{i-1}) > \rho(S_i) < \rho(S_{i+1}). \end{array}$$

It is then guaranteed that every cut piece has length at most 4, since without an equal-neighbor or extremum cut, a longer word would require a strictly monotone sequence of 5 ranks. Together with the retained exact intervals, these cut pieces yield

$$\mathsf{A}(S) = (z_0, \ldots, z_{v-1}), \qquad 0 = b_0 < \cdots < b_v = n, \qquad \|z_i\| = S[b_i : b_{i+1}).$$

**Algorithm 3** FillGaps: complete the exact partition and encode its pieces

**Require:** $S$, $n = |S|$, and ordered disjoint records $\mathcal{C}'$ from Resolve
**Ensure:** Exact pieces $\mathsf{A} = (z_0, \dots, z_{v-1})$ and boundaries $\mathbf{b} = (b_0, \dots, b_v)$

$(\rho(A), \rho(C), \rho(G), \rho(T)) \leftarrow (0, 1, 2, 3)$
$\mathcal{P} \leftarrow$ retained intervals interleaved with all gaps in $[0, n)$
$\mathcal{F} \leftarrow [\,]$
**for** each segment $[a, e)$ in $\mathcal{P}$ **do**
  **if** the segment is a retained record $(a, e, p, w)$ **then**
    Append $(a, e, w)$ to $\mathcal{F}$
  **else**
    Initialize cut$[a], \dots,$ cut$[e]$ to false
    cut$[a] \leftarrow$ true; cut$[e] \leftarrow$ true
    **for** $i = a, \dots, e-1$ **do**
      **if** $S_i = N$ **then**
        cut$[i] \leftarrow$ true; cut$[i+1] \leftarrow$ true
    **for** $i = a+1, \dots, e-1$ **do**
      **if** $S_{i-1} = S_i$ **then** cut$[i] \leftarrow$ true
    **for** $i = a+1, \dots, e-2$ **do**
      **if** $S_{i-1}, S_i, S_{i+1} \in \Sigma$ **then**
        **if** $\rho(S_{i-1}) < \rho(S_i) > \rho(S_{i+1})$ **then**
          cut$[i] \leftarrow$ true
        **else if** $\rho(S_{i-1}) > \rho(S_i) < \rho(S_{i+1})$ **then**
          cut$[i+1] \leftarrow$ true
    $c \leftarrow a$
    **for** $d = a+1, \dots, e$ **do**
      **if** cut$[d]$ **then**
        $x \leftarrow S[c : d)$
        $w \leftarrow N$ if $x = N$, otherwise the primitive root of $x$
        Append $(c, d, w)$ to $\mathcal{F}$; $c \leftarrow d$
$\mathsf{A} \leftarrow [\,]$; $\mathbf{b} \leftarrow [0]$
**for** each $(a, e, w)$ in $\mathcal{F}$ **do**
  **if** $w = N$ **then**
    $z \leftarrow (N, e-a, 0, [\,])$ ▷ literal ambiguity tile
  **else**
    $z \leftarrow (\sigma(w), e-a, \phi(w), [\,])$ ▷ identity, span, phase, empty script
  Append $z$ to $\mathsf{A}$ and $e$ to $\mathbf{b}$
**return** $(\mathsf{A}, \mathbf{b})$

**Canonical gap witnesses.** The proposal step (line 12 of Algorithm 1) uses the deterministic procedure $\Gamma$ below to explain fragments adjoining or separating anchors. A *gap context* is $C = (x, w_L, w_R)$, with at least one endpoint word specified, where $w_L$ fixes the template's starting phase, and the $w_R$ the phase immediately after it. We set

$$C^\dagger = (x^\dagger, w_R^\dagger, w_L^\dagger), \qquad \widehat{C} = \min\{C, C^\dagger\}, \qquad \perp^\dagger = \perp,$$

using a fixed lexicographic order on observed words. For a cap $0 \le d \le K$, we define a directional witness $g_d(C)$ by setting $g = |x|$ and enumerating integer lengths

$$\max(0, g-d) \le \ell \le g+d, \qquad w(\ell) = \begin{cases} w_L, & w_L \ne \perp, \\ \Phi_{-\ell}(w_R), & w_L = \perp. \end{cases}$$

We insert $x_\ell \cdots x_{g-1}$ at template offset $\ell$ when $g > \ell$, or delete the terminal $\ell - g$ template bases when $\ell > g$. Its elementary cost is

$$e_C(\ell) = |g - \ell| + \sum_{j=0}^{\min(g,\ell)-1} \mathbf{1}\{x_j \neq w(\ell)_{j \bmod p}\}, \qquad p = |w(\ell)|.$$

Among lengths with $e_C(\ell) \leq d$, we select the smallest tuple $(e_C(\ell), |g - \ell|, \ell)$ and return $\perp$ otherwise. To return a witness in the original orientation, we apply the reverse-complement mirror, which replaces the unit by $\sigma^\dagger$ and the phase by $-(\ell + \varphi) \bmod |\sigma|$, and reverses the script list. Consequently, an operation at template offset $z$ consuming $c$ template bases moves to $\ell - z - c$; emitted bases are complemented and deletion lengths are preserved. With $\perp^\dagger = \perp$, we define

$$\mathit{\Gamma}_d(C) = \begin{cases} g_d(\widehat{C}), & C < C^\dagger, \\ \big(g_d(\widehat{C})\big)^\dagger, & C > C^\dagger, \\ g_d(C), & C = C^\dagger \text{ and } g_d(C) = \big(g_d(C)\big)^\dagger, \\ \perp, & \text{otherwise.} \end{cases} \tag{1}$$

**Generating extension and join proposals.** Algorithm 5 implements line 12 of Algorithm 1 over $S$ and the array $\mathcal{C}$ of per-period interval lists, with the class rank, cap $k$, and budget table fixed for the construction. A proposal $q = (a; t)$ stores a raw-piece start index and its replacement tile with observed interval $I_q = [b_a, b_a + s_t)$. Since every proposal ends on a raw-piece boundary there is a unique index $r_q > a$ satisfying $b_{r_q} = b_a + s_t$ determined by the span. The proposal replaces $z_a, \ldots, z_{r_q - 1}$ by $t$; two proposals overlap in raw indices exactly when their observed intervals overlap. For anchors $l \leq r$ of the same identity and raw endpoints $a \leq l \leq r < b$, we assemble

$$\mathit{\Gamma}(S[b_a : b_l), \perp, w_l) \odot z_l \odot \big(\mathit{\Gamma}(S[b_{l+1} : b_r), \mathit{\Phi}_{s_l}(w_l), w_r) \odot z_r\big) \odot \mathit{\Gamma}(S[b_{r+1} : b_b), \mathit{\Phi}_{s_r}(w_r), \perp).$$

where the operator $\odot$ adds observed and template lengths and shifts each script's offsets by the preceding template length. We accept the assembled tile $t = (\sigma, s, \varphi, \mathcal{X})$ only if

$$\ell_t \geq 2|\sigma|, \qquad e_t \leq \beta_k\big(\mathrm{W}(\sigma, \varphi, \ell_t)\big). \tag{2}$$

**Algorithm 4** Propose: assemble and certify a periodic replacement

**Require:** $S$, raw pieces $\mathsf{A}$, boundaries $\mathsf{b}$, cap $k$, and budget table $\beta_k$
**Require:** $0 \leq a \leq l \leq r < b \leq v$; with $l, r$ with the same identity
**Ensure:** An accepted proposal $(a; t)$ or $\perp$

```
function Propose(a, l, r, b)
    f_L ← Γ_k(S[b_a : b_l), ⊥, w_l)
    if f_L = ⊥ then return ⊥
    t ← f_L ⊙ z_l; k' ← k − e(f_L)
    if l < r then
        f_M ← Γ_{k'}(S[b_{l+1} : b_r), Φ_{s_l}(w_l), w_r)
        if f_M = ⊥ then return ⊥
        t ← t ⊙ f_M ⊙ z_r; k' ← k' − e(f_M)
    f_R ← Γ_{k'}(S[b_{r+1} : b_b), Φ_{s_r}(w_r), ⊥)
    if f_R = ⊥ then return ⊥
    t ← t ⊙ f_R; compute its template length ℓ_t and edit cost e_t
    if ℓ_t < 2p_l or e_t > β_k(W(σ_t, φ_t, ℓ_t)) then return ⊥
    return (a; t)
```

We enumerate proposals in two passes:

1. At each anchor $i$, FreeExtensions tests each successive one-sided extension, adding at most $4p_i$ bases and stopping at the first failure and retaining successful extensions.
2. From each right anchor $r$, LeftAnchor scans left to the first anchor $l$ of the same identity, allowing at most $4p_r$ intervening bases, stopping at an incompatible anchor of span $> p_r + 1$ and returning $\perp$ if no anchor is reached. The four combinations $a \in \{l, A_l\}$ and $b \in \{r+1, B_r\}$ is tested for each selected pair.

Finally, we stably sort accepted proposals by $(a, s_t, e_t)$ using counting passes, and retain the first for each start-span pair $(a, s_t)$.

**Algorithm 5** Generate: extend anchors and join nearby anchors

**Require:** $S$, array $\mathcal{C}$ of per-period interval lists; fixed class rank, cap $k$, and budget table
**Ensure:** $(\mathcal{Q}, \mathsf{A}, \mathbf{b})$, with at most one proposal per start-and-span pair

```
C' ← Resolve(S, C)                                        ▷ Algorithm 2
(A, b) ← FillGaps(S, C')                                  ▷ Algorithm 3
v ← |A|; Q ← [ ]
for each anchor i do
    (A_i, B_i) ← (i, i+1)
    for d ∈ {L, R} do
        for (a, b) in FreeExtensions(i, d, 4p_i) do
            q ← Propose(a, i, i, b)                       ▷ Algorithm 4
            if q = ⊥ then break
            Append q to Q
            Update A_i ← a or B_i ← b in direction d
    if A_i < i and B_i > i+1 then append Propose(A_i, i, i, B_i) to Q
for each right anchor r do
    l ← LeftAnchor(r, 4p_r)
    if l ≠ ⊥ then
        for a ∈ {l, A_l} and b ∈ {r+1, B_r} do append Propose(a, l, r, b) to Q
Sort Q by (a, s_t, e_t)
Retain first for each (a, s_t)
return (Q, A, b)
```

**Selecting and emitting tiles.** Algorithm 1, lines 14–27, selects and emits proposals. Let $H$ be a 64-bit deterministic hash function, and let

$$H_\dagger(x) = \min\{H(x), H(x^\dagger)\}, \qquad \kappa(q) = (-\ell_t, e_t, H_\dagger(x_q)).$$

The surviving set is

$$\mathcal{V}(S) = \{q \in \mathcal{Q}(S) : \kappa(q) < \kappa(q') \text{ for every } q' \neq q \text{ in } \mathcal{Q}(S) \text{ with } q \sim q'\}. \tag{3}$$

Every comparison uses the original proposal set, and equal and overlapping keys both lose even when equality results from a hash collision. Consequently, survivors are disjoint. We emit a survivor's tile and skip its support, or emit the next unchanged raw piece in order, which yields $\tau_k(S)$.

**Construction cost.** Assume a word RAM with fixed-size hashes and $\mathcal{O}(1)$ lookup. The notation $O_P(\cdot)$ permits constants depending on $P$.

**Proposition 3 (Construction time and space).** *For nonempty input, construction takes $O_P(n(k+1))$ time and working words. For fixed supported cap $K$, working space is linear in $n$.*

*Proof.* The $P$ scans and merges take $O_P(n)$ time. Each merged interval requires processing at most $P$ heads, and recovering its initial word at most $P$ bases. By Fine–Wilf, distinct maximal primitive runs of periods $p, q \le P$ overlap in fewer than $p + q - \gcd(p, q) < 2P$ bases. Removing contained intervals, each interval has at most $2P$ overlapping intervals. All constraints and cuts therefore take $O_P(n)$ time, producing $v \le n$ pieces.

Each anchor makes $O_P(1)$ extension and bridge attempts, including failures, with at most three gaps of length at most $4P$. A gap with remaining cap $d \le k$ tests at most $2k + 1$ template lengths, comparing at most $4P$ bases per length. Stable counting passes, constructing or mirroring scripts, and concatenating fragments all take $O_P(n(k + 1))$ time and space.

Hashing proposals also has a linear aggregate bound. Since endpoint pairs are unique and widths are at most $W_0$, each raw piece belongs to at most $\sum_{h=1}^{W_0} h$ proposals. Consequently,

$$\sum_{q \in \mathcal{Q}(S)} |x_q| = \sum_{i=0}^{v-1} s_i \,\#\{q : a_q \le i < r_q\} \le \frac{W_0(W_0 + 1)}{2}\, n. \tag{4}$$

Computing and caching both orientations of each hash therefore costs $O_P(n)$, including proposals with long anchors or colliding hashes. At most $W_0$ deduplicated proposals start at any index, so the survivor test compares $O_P(1)$ cached keys; advancing past covered pieces costs $O(v)$ in total. Summing these bounds proves the claim.

##### A.3 Compatibility with sequence operations

We establish consistency under reverse complementation, restriction to a substring, and a single nucleotide edit. Throughout, $P \ge 4$ and $0 \le k \le K$ are fixed, and $\tau_k$ is the bounded construction of Subsection A.2.

**Theorem 4 (direct mirror).** *Let $t = (u, s, \phi, \mathcal{X})$ have a valid ordered script, period $p = |u|$, and template length $\ell = s + D - I$. Reversing the exception list moves an operation at template offset $z$ that consumes $c$ template bases to offset $z' = \ell - z - c$, where $c = 0$ for an insertion, $c = 1$ for a substitution, and $c$ is the deleted run length for a deletion. Then*

$$\left\|t^\dagger\right\| = \|t\|^\dagger, \qquad (t^\dagger)^\dagger = t.$$

*Proof.* For $0 \le j < \ell$, the mirrored template has nucleotide

$$W(u^\dagger, -(\ell + \phi), \ell)_j = \overline{u_{(\phi+\ell-1-j) \bmod p}} = \big(W(u, \phi, \ell)^\dagger\big)_j,$$

with phases read modulo $p$. Interval $[z, z + c)$ is positionally mapped to $[\ell - z - c, \ell - z)$, and an insertion at boundary $z$ moves to boundary $\ell - z$. Valid script order is preserved in the reverse list: $z_i + c_i \le z_{i+1}$ becomes the corresponding ordered nonoverlap condition after reflection. The transformed script therefore renders the reverse-complemented observation. Applying it twice restores offsets and phases, since

$$\ell - (\ell - z - c) - c = z, \qquad -\big(\ell - (\ell + \phi)\big) \equiv \phi \pmod p.$$

**Proposition 4 (construction symmetry).** *For a sequence $S$ of length $n$, reflection of nucleotide intervals $[a, b) \mapsto [n - b, n - a)$ sends its raw partition to that of $S^\dagger$. If $v = |\mathsf{A}(S)|$ and $q = (a; t)$ has end index $r_q$, the map*

$$(a; t) \longmapsto (v - r_q; t^\dagger)$$

*is a bijection between the accepted proposal sets $\mathcal{Q}(S)$ and $\mathcal{Q}(S^\dagger)$.*

**Theorem 5 (Independent strand equivariance).** *For every input sequence $S$,*

$$\tau_k(S^\dagger) = \big(\tau_k(S)\big)^\dagger.$$

*Proof.* Proposition 4 preserves overlaps, template lengths, costs, and the strand-neutral content hash. Hence, it preserves the priority key and every strict comparison in (3).
